# Evaluating codon optimization strategies for mammalian glycoprotein production with an open-source expression vector

**DOI:** 10.64898/2026.03.18.712111

**Authors:** Chang Yang, Raina Soni, Sarah E. Visconti, Mina Abdollahi, Filmawit Belay, Anita Ghosh, Samuel W. Duvall, Curtis J. W. Walton, Rob Meijers, Haisun Zhu

**Author notes:** Correspondence should be addressed to CJWW, RM and HZ.

## Abstract

Efficient production of human proteins for the development of tool compounds and biologics depends on a detailed understanding of the protein expression machinery in mammalian cells. Codon optimization is widely believed to enhance protein yield, yet its impact in homologous mammalian systems remains poorly defined. Here, we systematically compare five codon usage strategies reflecting common assumptions about rare codons, RNA stability, and synthesis efficiency. We developed pTipi, an efficient open-source mammalian expression vector, and evaluated its performance in antibody production. We generated plasmids for common epitope tag antibodies such as V5, anti-biotin and anti-His for distribution by Addgene. To compare codon usage schemes, we performed a bake-off of 18 human and murine Wnt pathway glycoproteins in mammalian cells. Small-scale expression screens revealed that codon optimization did not provide a general advantage over native coding sequences, while strategies prioritizing RNA stability consistently reduced expression. Interestingly, a skewed codon scheme using the most abundant codons produced yields comparable to native sequences and occasionally enhanced protein output. To enable flexible evaluation of codon strategies, we implemented a Golden Gate–compatible pTipi platform for efficient synthetic gene incorporation. We conclude that native codons are sufficient for robust homologous mammalian expression of glycoproteins, while selective codon skewing can be beneficial for some targets.

## Introduction

The production of human glycoproteins in mammalian cell lines has been well established for decades, with a myriad of synthetic tools available to boost protein yield^1^. Careful construct design is an important factor in obtaining high expression yields. While machine learning (ML) methods have made strides in defining construct boundaries^2^, other factors that are important in efficient protein production remain poorly understood. One important factor is the regulation of protein expression through the redundancy of the genetic code, where multiple codons code for the same amino acid^3^. Codon usage affects DNA transcription^4^, RNA translation and stability^5^, as well as many other regulatory mechanisms including interactions with non coding RNA^6^, the spliceosome^7^ and DNA and RNA binding proteins^8^. Codon optimization attempts to boost expression by addressing the liabilities that apply to all genes, including the use of rare codons, RNA stability and GC content. The codon adaptation index (CAI) measures how closely the codon usage of a gene matches the codon usage bias of a reference set of highly expressed genes in the expression host^9^. It is commonly used to guide optimization of heterologous protein expression. CAI values range from 0 to 1, where a value of 1 indicates exclusive use of the most preferred codons in the host reference set. More recently, ML tools have become available to improve RNA stability, as RNA half-life can improve protein expression levels^10^.

Systematic studies on the impact of codon usage on protein expression have been performed for *E. coli*^11^, but no such systematic study has been done for mammalian expression systems to our knowledge. Nevertheless, codon optimization is widespread in the biotechnology industry, despite a lack of understanding of its effect on mammalian protein expression^12^. There is anecdotal evidence the expression of individual proteins benefitted from codon modifications, for heterologous expression of human proteins in prokaryotes, and viral proteins in mammalian cells^13^. It is believed that some human protein families are strongly regulated on the transcription and translational level due to their potency in cell signaling^14^. The Wnt pathway for instance affects ribosomal translation regulation^15^, indicating that an underlying feedback mechanism may adversely affect overexpression of Wnt receptors. This behavior suggests the expression of these human gene products in HEK cells may benefit from codon optimization.

Here, we performed a systematic comparison of codon optimization schemes for several members of the Wnt receptor^16^ and carrier^17^ families, testing both the human and murine orthologs. To enable this comparison, we generated a minimal expression vector (pTipi) containing only those components considered essential for transient mammalian protein expression. We performed an initial small scale expression campaign to compare expression levels for five codon usage schemes, followed by comparative large scale expression and purification for several of the glycoproteins.

These results indicate that native codon usage is remarkably effective compared to codon optimized sequences and is never detrimental. Strategies that focus on RNA stability consistently led to low protein expression yields, whereas a codon scheme that only uses the most abundant codons (“skewed codons”) occasionally increased expression compared to a native codon scheme. Based on these observations, we implemented a Golden Gate cloning scheme for the pTipi vector that enables the efficient incorporation of inserts representing different codon schemes. These experiments inform an overall strategy for the application of codon usage strategies in high throughput mammalian protein production that is widely applicable.

## Results

### Design and testing of an open-source transient mammalian expression vector

We designed the minimal pTipi2.1 expression vector (Addgene #229144) to incorporate essential, off-patent elements for transient expression of glycoproteins, while minimizing the size of the vector by removing stuffer sequences between the elements (Figure 1A). The pUC18 plasmid was used as a backbone^18^, and the original lacZ expression cassette was replaced with a customized DNA fragment for mammalian expression. This fragment consisted of a CMV enhancer and promoter cassette^19^, immediately followed by a synthetic chimeric intron^20^, a Kozak sequence^21^, a human Azurocidin signal peptide^22^, a multiple cloning region to insert the gene of interest, followed by the Woodchuck Posttranscriptional Regulatory Element (WPRE)^23^ and an HSV poly adenylation signal sequence^24^. The vector is used for transient protein expression in common mammalian cell lines such as HEK293 and CHO, and is considerably smaller in size than most commercially available vectors.

**Figure 1.**
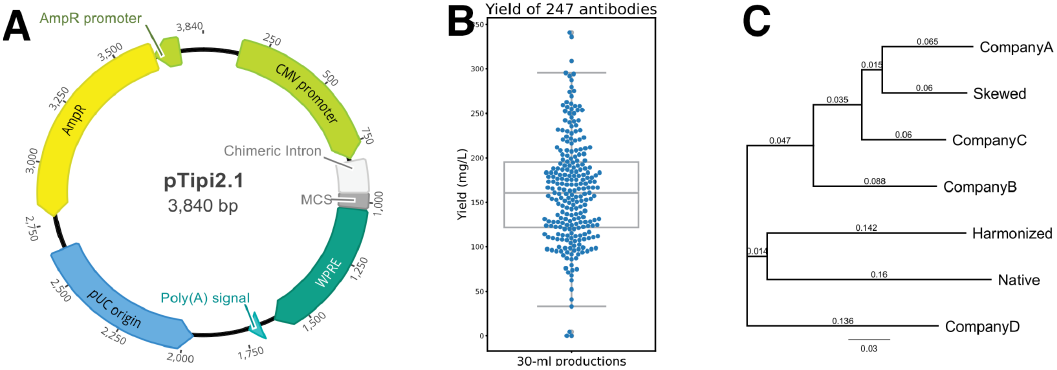
Open-source pTipi vector design and performance. **(A)** vector map of the pTipi vector showing the different elements and restriction sites that can be used for restriction site cloning to insert a fragment after the chimeric intron, with or without the signal peptide from human Azurocidin. **(B)** Protein yields for antibodies produced using the pTipi2.1 expression vector after one step of affinity purification with protein A. **(C)** Dendogram of the different codon schemes obtained for a human LRP6 construct, showing relationships with the native codon scheme and a skewed codon scheme that only uses the most abundant codon for each amino acid.

To show the utility of the pTipi2.1 vector, we tested expression of popular epitope tag antibodies in this vector. We fused the published sequences of the variable domains of Anti-Biotin (M33), Anti-EE (3D5EE_48.K)^25^, anti-GCN4 (C11L34)^26^, anti-Strep (C23.21), anti-Flag (4E11/M1)^27^, anti-V5 (SV5-Pk1)^28^, anti-ProC (HPC-4)^29^, anti-His (HAP1/5.13.22)^30^, anti-Rho (Rho 1D4)^31^ to the constant domains of rabbit IgG. We performed cotransfection in

Expi293F cells with the heavy and light chain each incorporated into separate pTipi2.1 vectors, at a heavy/light chain plasmid ratio of 1:2.5. The chimeric antibodies were purified from the culture soups using protein A resin. The protein yields ranged from 30 to 160 mg/liter, except for the anti-V5 antibody which had a relatively low yield of 13.8 mg/L (Table S1). The chimeric rabbit isotype IgG antibodies and pTipi2.1-based heavy/light chain plasmid pairs are available through [Addgene](https://www.addgene.org/antibodies/tags-markers/) for general academic and commercial use.

Since the chimeric nature of these antibodies may negatively affect expression yields, we also expressed a number of standardized reagent antibodies developed via our discovery platform, using the same coexpression approach for the heavy and light chain antibody plasmids. The median yield of purified protein is 160.7 mg/L for 247 antibodies (Figure 1B). These are typical yields for antibodies produced by transient expression in mammalian cells, showing the pTipi2.1 vector is a reliable resource for transient mammalian glycoprotein production.

### A codon usage bake-off experiment

In the process of ordering a batch of synthetic genes related to the Wnt pathway family, we obtained codon optimization proposals from four synthetic gene providers and noticed some remarkable trends in codon usage. The algorithms of several vendors suggested codon optimized genes that skewed codon usage of amino acids with degenerate codes like leucine, arginine and serine towards the more abundant codons. To illustrate the issue, we made a comparison in codon usage for a construct of LRP6. When analyzing leucine codon usage, all four vendors avoided the rare CTA codon, and three vendors favored the most frequent CTG codon for the vast majority of leucine codons (Table S2). A phylogenetic tree of the nucleotide sequences for the LRP6 construct shows that Companies A-C are related to a design that only uses the most abundant codons (Figure 1A). We called this approach to codon optimization the “Skewed” method, and decided to investigate how it compared to other codon usage methods in a bake off experiment involving 9 human glycoproteins and their murine orthologs. We selected glycoproteins involved in the Wnt signaling pathway, creating truncated constructs of domains that are directly involved in binding Wnt (Figure 2), with the ultimate aim to create antibodies that may interfere with Wnt binding. The sequence identity between the human and murine homologues is 93 % or higher, which creates the opportunity to study the effect of codon optimization schemes on closely related genes (Table S3). We created two truncated ectodomain constructs for LRP5 and LRP6 covering adjacent betapropellor domains for comparison.

**Figure 2.**
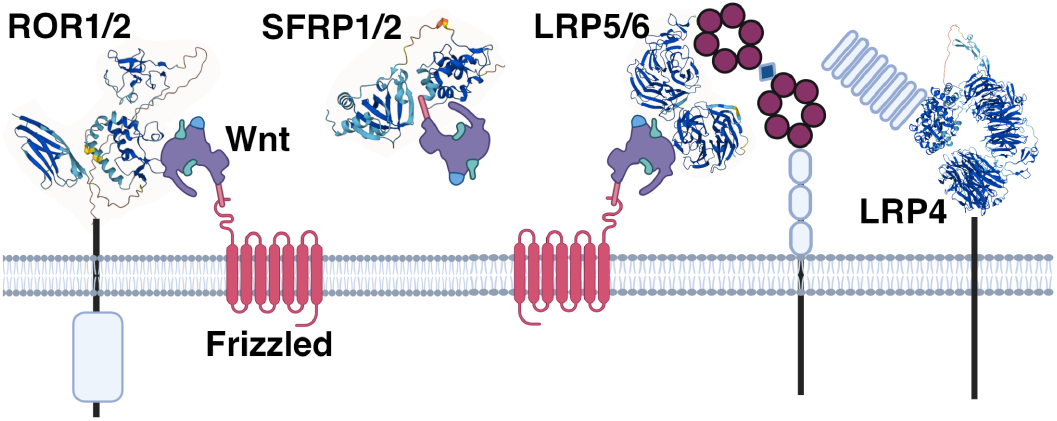
Wnt receptor glycoproteins tested. Human Wnt receptors tested with different codon schemes. Constructs are depicted as ribbon diagrams for ROR1, ROR2, LRP4, LRP5 and LRP6 ectodomains and the secreted glycoproteins SFPR1 and SFPR2. Schematics of the Wnt ligand and Frizzled receptors (Fzd) are shown to indicate the mode of interaction. The picture was produced with BioRender.

We designed three codon usage scenarios that would emphasize different philosophies for codon optimization: 1) the use of abundant codons (Skewing method), 2) codon redistribution (the Harmonization method), and 3) optimize RNA stability (the LinearDesign method). For the Skewing method, we used the single most abundant codon for each amino acid. For the Harmonization method, we redistributed the rare codons using the Charming software^32^, maintaining a CAI index that is similar to the native DNA sequence. For the LinearDesign method, we used the LinearDesign algorithm to optimize the free energy of the predicted RNA structure, while keeping the CAI index of optimized sequences above 0.8 to ensure that expression is not affected by an abundance of rare codons. For comparison, we also created constructs identical to the native DNA sequences, and we also considered codon optimized genes from two of the companies (Company A and D) using their proprietary algorithm. Only the gene insert was codon optimized, while the signal peptide and purification or identification tags all had the same nucleotide sequence.

Analysis of the overall codon usage by each codon optimization strategy showed that Native and Harmonized codon usage had similar CAI indices, GC content and codon frequencies, as expected (Figure 3 and Supplementary Figure S1). The Native codon usage was based on cDNA derived from the Refseq database^33^, and the overall codon usage for these constructs closely matched average codon usage for the human and mouse genome (Figure 3A).This shows that the glycoproteins chosen for this comparison have a codon distribution that is typical for human genes, and there are no anomalies. The Skewed codon design used the single most abundant codon across all targets and consequently had the highest average GC content (Supplementary Figure S1) as well as maximum CAI score of 1. LinearDesign sequences had a slightly elevated GC content compared to the Native design. Company A showed persistent skewing of the more degenerate codons towards high frequency codons, but maintained an average GC content similar to the Native sequence (Figure 3A). The method employed by Company D consistently reduced the usage of the dominant codon for each amino acid while avoiding the use of one Leucine (CTA) and one Serine codon (TCG) altogether.

**Figure 3.**
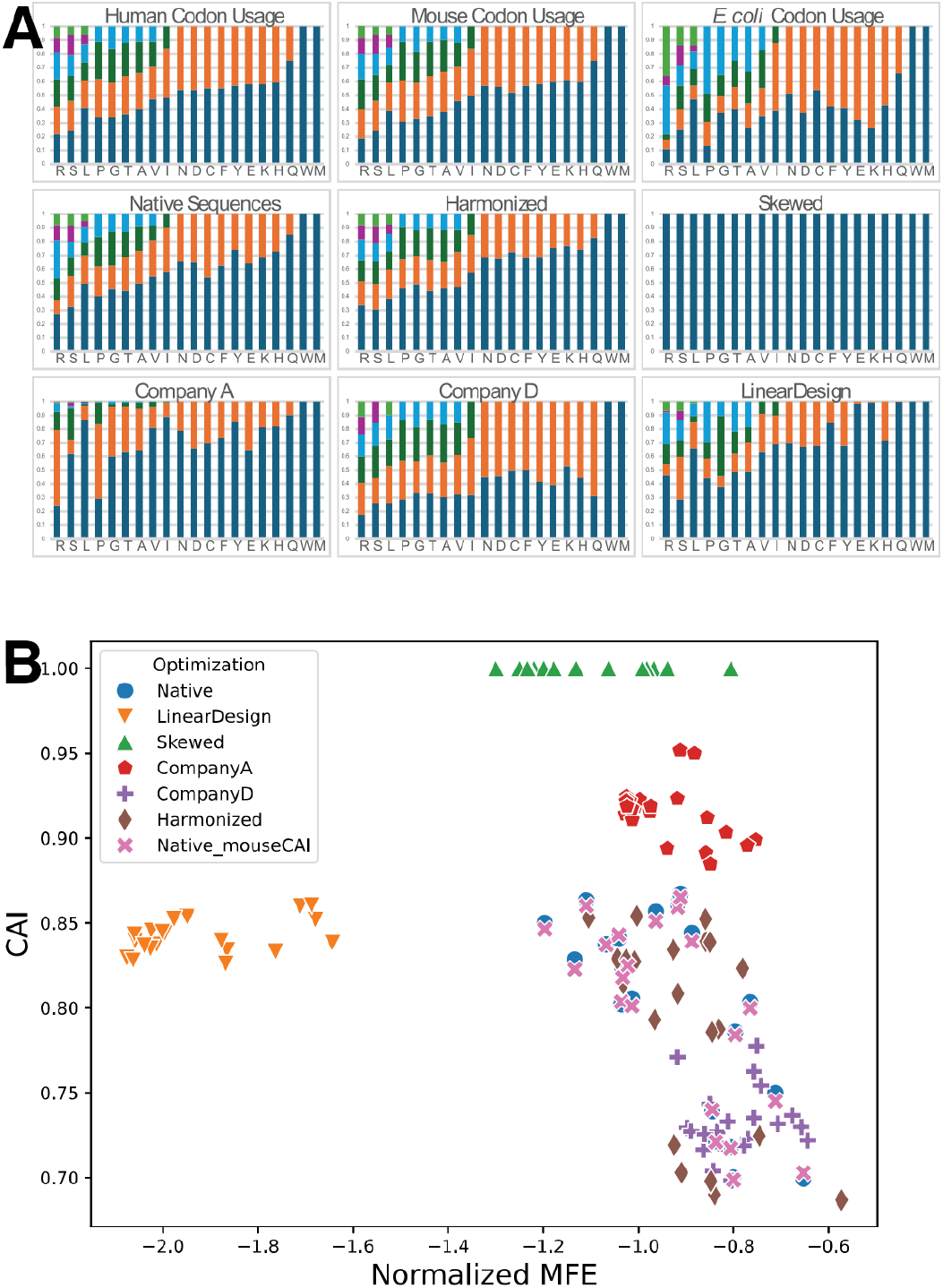
Comparison of codon usage among optimization schemes. **(A)** For each optimization scheme, the average codon usage frequencies for all 20 amino acids were calculated from 18 sequences and plotted in a stacked bar chart. The average codon usage for the native human, murine, and *E. coli* DNA sequences were plotted as reference. For each amino acid, the order of the codon was based on the frequency in human codon usage, with the most abundant at the bottom and the least abundant at the top. **(B)** Mapping rare codon usage versus RNA stability for the different codon usage schemes. Graph showing the mean free energy (MFE) normalized by amino acid length of the construct versus CAI index. Each construct is colored according to its codon optimization scheme.

The most important factors considered in codon optimization are rare codon distribution and mRNA stability, so we plotted the CAI index versus the minimum free energy (MFE) of the mRNA, normalized by sequence length (Figure 3B). The native codon scheme covered a narrow range, with a CAI index lying between 0.7 and 0.87. For reference, we also calculate the CAI scores of native sequences based on mouse codon usage. Because human and mouse genes share highly similar codon usage patterns (Figure 3A), the CAI scores for each construct were almost identical. Interestingly, the range for the normalized MFE is also relatively narrow for the native codon scheme, especially when compared to the LinearDesign scheme which focused on optimizing mRNA stability. The normalized MFE is two-fold lower for LinearDesign compared to the native nucleotide sequence, while both have the same spread of approximately 0.5. The Harmonized scheme closely tracked native codon usage, as it only more evenly distributed codons from the original Native codon sequence. The Skewed scheme showed a slightly lower normalized MFE than the Native codon scheme, although this was not considered in choosing codons, indicating that the more common codons also may contribute to a more stable mRNA structure. For comparison, the codon usage schemes obtained from algorithms from Company A and D each cluster together. Company A’s scheme had a relatively high CAI index range (from 0.85 to 0.95) with a normalized MFE that was similar to the native codon scheme. Company D’s scheme showed a lower range for the CAI index (0.68 to 0.80) and had the highest average normalized MFE of all codon usage schemes. We decided to include the codon optimized sequences from Company D in a comparative expression screen *bake-off*, as these constructs had the lowest average CAI index, the highest normalized MFE, and the most evenly distributed codon selection.

### Performance of codon usage schemes in small scale expression

To compare the expression levels of the different codon usage schemes, we performed a small scale protein expression screen^34^. All the constructs were equipped with an Avi tag for biotinylation by Biotin ligase and an 8x His tag at the C-terminus, separated by GSG linkers. The protein constructs were expressed in Expi293F cells in a 384 deep well plate, and biotinylated in media after removal of the cell pellet. Protein levels were detected using RDye^®^800CW Streptavidin and compared to protein standard. Expression could be detected for all constructs, except for the LinearDesign scheme of murine LRP4. The expression screen was performed in triplicates, and some general trends can be discerned in the performance of the codon usage schemes. The most prominent trend is that the LinearDesign method which emphasizes RNA stability was the poorest performer overall, and often showed significantly lower expression levels than the other codon usage schemes (Figure 4A and 4B). It appeared that the Harmonized codon scheme gave the most variable expression, and it performed worse when used on murine ortholog constructs (Figure 4B). The comparative performance of the other codon usage schemes was more subtle. We ranked the expression levels of the five codon schemes for each construct to compare overall performance (Figure 4C and Supplementary Figure S2). Interestingly, while Harmonized was the best performing codon scheme for 8 of the 18 constructs, it was also the worst performer for 7 constructs among the productive codon schemes (Native, Harmonized, Skewed). The Skewed codon scheme did relatively well, and was the best performer for 4 of the 18 constructs. However, 7 out of 18 Skewed codon constructs performed worse than other productive codon schemes. It was somewhat surprising that the radical skewing of the codon usage was not detrimental to small scale expression. To conclude, it appeared that codon schemes that address issues around codon distribution all gave similar expression levels comparable to the native codon scheme, but the codon scheme that emphasized RNA stability was detrimental.

**Figure 4.**
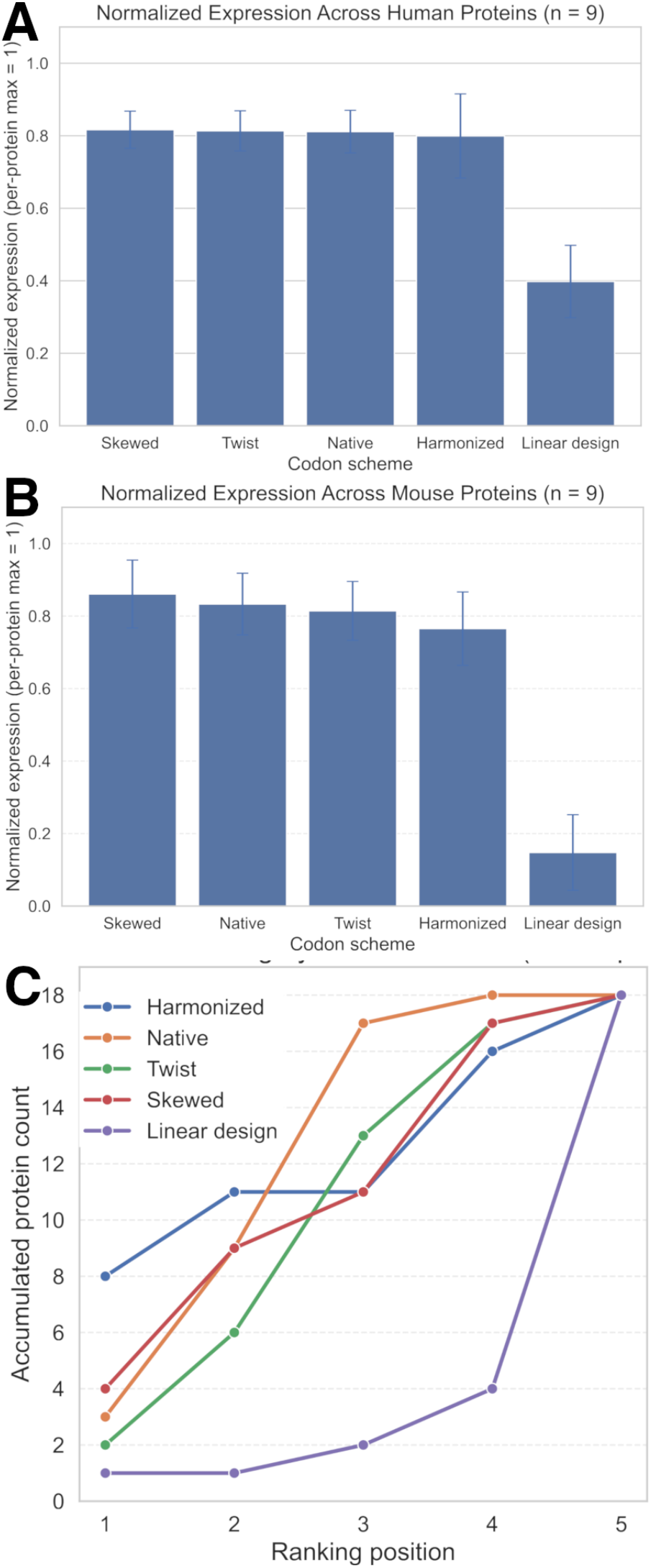
Small scale expression performance for 18 glycoproteins using 5 codon schemes. **(A)** Normalized expression across human proteins (n = 9) for each codon scheme. For each protein, expression values were normalized to the highest-expressing scheme (set to 1.0). Bars show the mean normalized expression across proteins; error bars denote the standard error of the mean (SEM) across proteins. **(B)** Normalized expression across mouse proteins (n = 9), calculated. **(C)** Accumulated ranking analysis of codon scheme performance across all 18 proteins. Codon schemes were ranked per protein based on normalized expression (rank 1 = highest). Curves indicate the cumulative number of proteins for which a given scheme achieved a given rank or better. The native codon scheme accumulates top rankings most rapidly and reaches the full protein set at earlier ranking positions, indicating the most consistent overall performance across targets.

### Large scale production and validation of ROR1, ROR2 and LRP6 constructs

To verify the outcome of the small scale expression screen, we performed large scale (100 mL Expi HEK293 cell culture volume) expressions for human ROR1, ROR2 and murine LRP6 for all five codon schemes followed by affinity chromatography and size exclusion chromatography (SEC). The large scale expression screen confirmed that the LinearDesign method substantially lowered protein yield, and in case of LRP6, only traces of the protein could be detected (Figure 5). The other codon schemes gave similar yields with some variation, based on area under the curve calculations from the SEC runs (Table S4). Interestingly, the Skewed codon scheme gave the largest variation, emerging as the best performer for ROR1 and the worst performer for ROR2 among the 5 codon schemes. The glycoproteins produced are of high purity, showing a single prominent band on an SDS-PAGE gel (Figure 5). The proteins are monodisperse as observed by dynamic light scattering analysis, and do not aggregate after repeated freeze/thaw cycles.

**Figure 5.**
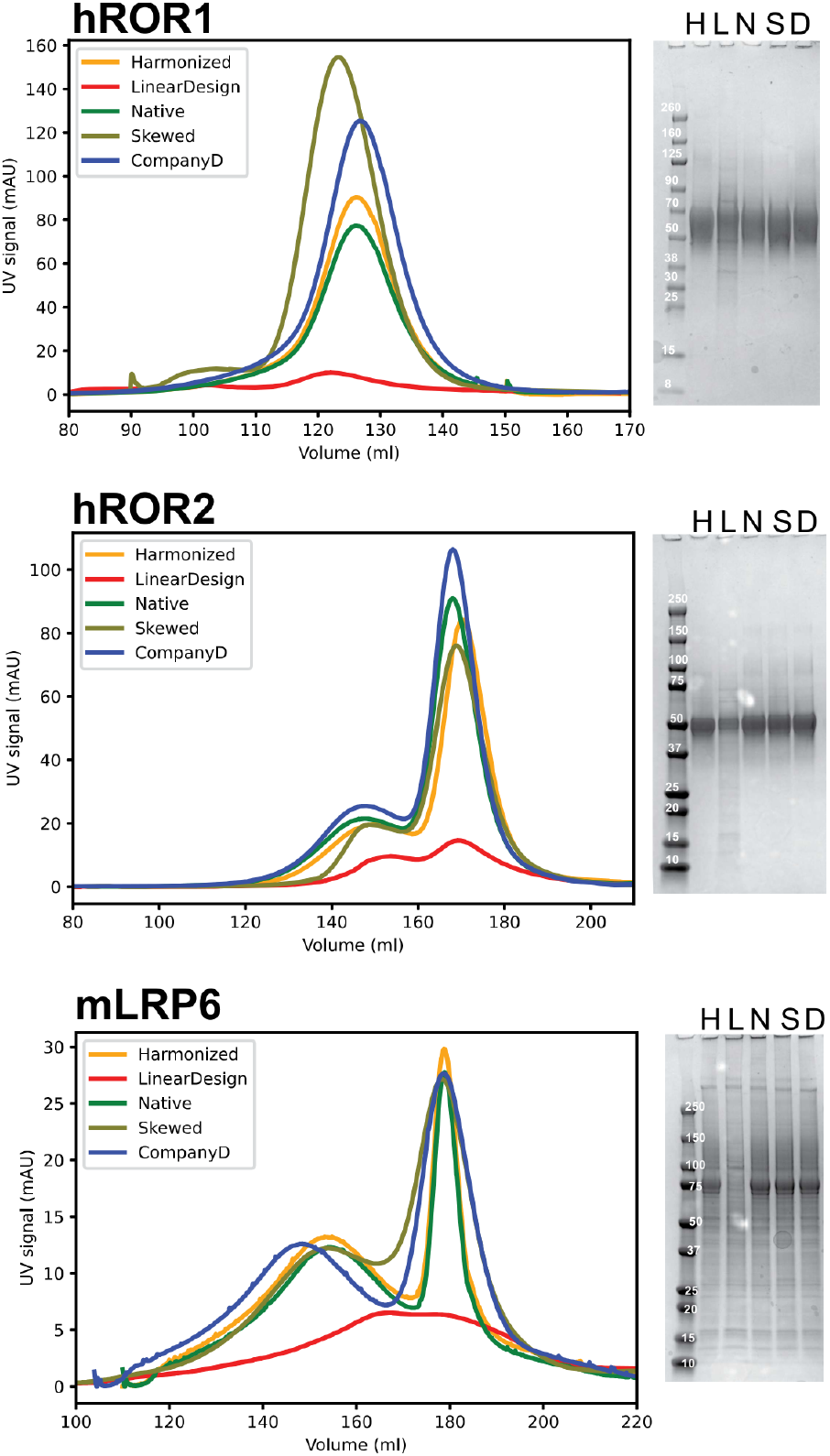
Comparative large scale production of three constructs using five codon schemes. Size exclusion profiles (left panels) and non-reduced SDS-Page gels for human ROR1 and ROR2 and murine LRP6_629-1244. H: Harmonized; L: LinearDesign; N: Native; S: Skewed; D: Company D

### Golden Gate assembly scheme for versatile synthetic gene incorporation

To enable the efficient incorporation of synthetic gene fragments with native codon schemes, we modified the pTipi expression vector to facilitate Golden Gate cloning^36,37^. The vector, pTipi2.2 (Figure 6A), was modified to accommodate the use of the BsaI, BsmBI and PaqCI type II restriction enzymes, which allows for scarless integration of a synthetic gene insert. The multiple cloning site was replaced with a GFP dropout cassette from pYTK047^38^, flanked by the Type IIS restriction enzyme site, PaqCI, and the backbone was cured of BsaI and BsmBI by introducing point mutations. To confirm the point mutations did not negatively affect the protein expression a comparison of protein yield between pTip2.1 and pTip2.2 was performed and no significant differences in protein yield were observed (Figure 6B). The pTipi2.2 vector is now routinely used to incorporate gene blocks for efficient protein expression.

**Figure 6.**
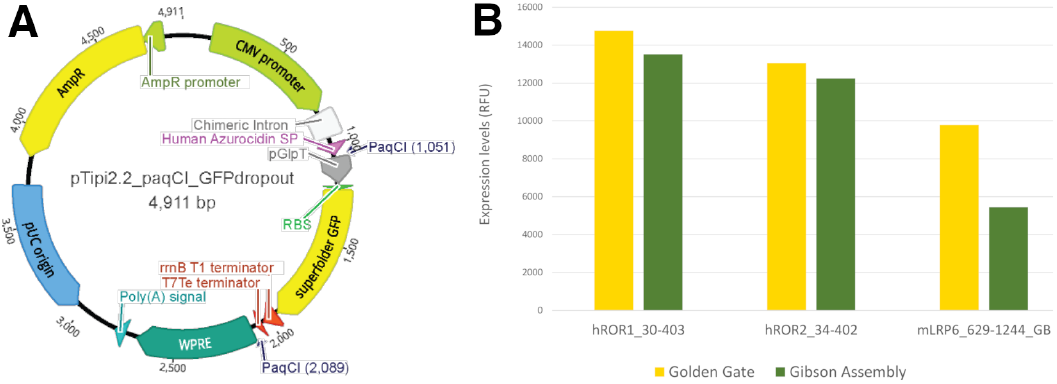
Golden gate conversion of pTipi expression vector. A) Plasmid map of the Golden gate cloning pTipi2.2 expression vector containing a GFP insert, with restriction sites for *BsmBI* and *PaqCI* removed. B) Comparison of small-scale expression yields when using Gibson cloning (pTipi2.1) or Golden Gate cloning (pTipi2.2) using identical inserts, to show the CMV promoter was not compromised after removal of the *BsmBI* restriction site.

## Discussion

The production of mammalian glycoproteins requires substantial efforts to obtain quality material at high concentrations^39^. In addition to overall yield, one must be mindful of simultaneously optimizing parameters to reduce costs and increase the success rate of glycoprotein production, including the development of cell lines and/or plasmids for transfection, expression conditions, purifications schemes and glycoprotein construct design. Construct design requires careful analysis of domain boundaries, the placement of tags and signal peptide for secretion and the use of fusion proteins to boost solubility and stability. Codon optimization has been another area of development, especially with the advances in gene synthesis which have made the cloning of truncated proteins into plasmids more affordable. The benefits of different codon schemes for mammalian expression have not been carefully evaluated to our knowledge, and this study aimed to compare widely used codon usage strategies and apply them to a set of challenging glycoproteins that serve as antigens in an antibody discovery campaign.

We first designed a minimal protein expression vector pTipi2.1 for transient expression in mammalian cells. We configured the vector down to the most essential elements, resulting in expression levels that are sufficient for small and large scale production of glycoprotein antigens and antibodies. This expression vector is readily available through Addgene with a non-restricted license for general academic and commercial use. We then designed constructs for members of the Wnt receptor and carrier families, with the rationale that these glycoproteins may benefit from codon optimization schemes. The different codon schemes we employed addressed elements of RNA translation, including RNA stability and tRNA processing. We found that none of the schemes systematically outperformed the other schemes or the use of the native nucleotide sequence, which indicates that codon optimization in homologous mammalian expression may not be necessary to improve protein production. This is in contrast with heterologous expression of human gene constructs in non-mammalian hosts^13^. Interestingly, we found that reducing the codon usage from 61 codons to the most commonly used 20 codons did not affect protein expression for most of our antigens.

Occasionally, the skewing of codon usage led to substantial improvement over native codon usage, indicating that the repetitive use of common codons may lead to efficient reuse of tRNAs^40^ and tRNA depletion is not an issue^41^. We conclude that the use of native codons is the strategy most likely to result in efficient protein production, but that secondary issues involving gene synthesis and removal of restriction sites might make alternate codon schemes attractive. In that case, it is important to evaluate the proposed codon usage scheme and extreme skewing of the codons may be a feasible alternative. To facilitate the autonomous and efficient cloning of synthetic genes with a preferred codon scheme, we modified the pTipi expression vector to enable Golden Gate cloning and made this vector available at Addgene. This reagent will empower labs to make their own decisions in the choice of codon usage for optimal protein expression.

## Acknowledgements

The pYTK047 plasmid was a gift from John Dueber (Addgene plasmid # 65154; http://n2t.net/addgene:65154; RRID:Addgene_65154). We would like to thank Meghan Rego and Steve Almo for reading the manuscript.

## Materials and Methods

### Constructing pTip2.1, a vector for mammalian protein expression

A novel plasmid vector, pTip2.1, was engineered for protein expression in mammalian cells. This vector’s foundation is the standard pUC18 plasmid, wherein the original lacZ expression cassette (431 bp) was substituted with a custom DNA fragment optimized for mammalian expression. This fragment integrates a standard CMV promoter, a chimeric intron derived from beta-globin and human IgG introns, a customized multiple cloning site (MCS), followed by the WPRE fragment and the HSV polyadenylation signal sequence. The resulting pTipi2.1 vector is relatively compact (3840 bp), which is anticipated to enhance cloning and protein production efficiency. This vector, along with a slightly modified variant incorporating the human Azurocidin signal peptide (pTipi2.1-AZU_SP), has been deposited with Addgene (Catalog #225877 and #229144) and are available for general use.

### Constructing pTip2.2, a Golden Gate compatible vector for efficient synthetic gene incorporation

To enable the rapid assessment of N-terminal and C-terminal tags on protein expression yield, the pTip2.1 vector was modified to be golden gate compatible using the high fidelity overhangs from Set 2 identified by Potapov et al ^42^. This set of overhangs will enable a high fidelity assembly and provide a sufficient number of additional overhangs for assembling large CDS parts ^43^ or expanding the partial library. The system is designed to allow for additional modularity to be introduced if the user desires swapping additional features on the backbone, such as the promoter, resistance markers and affinity or epitope tags. This vector has been deposited with Addgene (catalog #225877 and is available for general use.

### Assembling Expression Vectors

The gene or gene fragment of interest can be ordered or PCR amplified to include inward-facing paqCI restriction sites with Part 3 overhangs (see Table S5), The CDS part can also be flanked with BsaI sites for subcloning into the Entry vector for long term storage and amplification by *E. Coli*. The golden gate reaction is performed following the manufacturer’s recommended protocol (NEB) for the PacCI (NEB #R0745) and T4 DNA Ligase (NEB #M0202). Briefly, one of each part type (1-4) are mixed together using ~75 ng of each plasmid or 2:1 molar ratio of insert to backbone, 2 uL T4 Ligase buffer (10x), 5-10U of PaqCI (1uL), 200-400U of T4 Ligase (1 uL), 5 pmoles of PaqCI activator (0.25 uL), Water (to 20uL). Cycle the reaction from 37C, 1 min to 16C 1 min) for 30-60 cycles, heat inactivate at 60C for 5 minutes. The assembled reactions are chemically transformed following standard protocols, and plated on LB Agar Carbencillin or Ampicillin (100ug/mL) overnight at 37C. The colonies are checked for the absence of GFP using a blue light transilluminator. 2-3 white colonies are picked and sequence confirmed using whole plasmid sequencing.

### Sequence selection and codon optimization

#### Native sequences

For this study, 18 protein fragments from 16 genes were selected. Native DNA sequences were obtained from GenBank RefSeq records, corresponding to the chosen protein fragments. These sequences were subsequently cross-referenced with SwissProt/UniProt sequences. In two mouse sequences, mLRP6 and mROR2, specific amino acid discrepancies were observed between the GenBank and SwissProt entries. Since native DNA sequences are not available from SwissProt, the RefSeq sequences were utilized as templates for all subsequent optimization schemes.

#### Codon-skewed sequences

For the codon-skewed sequences, DNA sequences were reconstructed using the most frequent codons for each amino acid. Codon usage was determined by referencing a human codon usage table ^44^.

#### Harmonized sequences

We used the open-source Charming tool to redistribute the native codon sequence for codon harmonization ^32^. The built-in MinMax method was applied to evaluate a sliding window of 17 codons of each sequence. We used a reference codon frequency table to generate the optimized sequences ^44^.

#### LinearDesign sequences

LinearDesign is a software package that optimizes codons by considering both RNA minimal free energy (MFE) and the codon adaptation index (CAI) ^10^. The MFE is computed using the ViennaRNA method ^45^, while the CAI is derived from a standard codon usage table provided with the tool package. The software identifies sequences with the lowest MFE while maintaining a certaining level of CAI based on user input. Results from the original study showed that exclusively focusing on MFE did not consistently produce optimal expression yields. Consequently, we set a CAI threshold at 0.8 and selected the lowest free energy sequences generated by the software for this study.

#### Codon optimization by commercial vendors

Codon optimizations of the native sequences were performed using tools available from vendor websites (Company A-D). In all cases, the default settings were used and the only requirement was to optimize for expression in human cells with no restriction placed on potential enzymatic digest sites.

### Plasmid construction

All sequences, comprising both native and codon-optimized variants (Native, Skewed, Harmonized, LinearDesign, and Company D), underwent synthesis and subsequent cloning into the pTip2.1-AZU_SP plasmid at Genscript or Twist Biosciences. Each sequence additionally incorporates a fixed C-terminal AVI and HIS tag, facilitating purification and biotinylation.

### Small-scale protein expression and ELISA testing of expression level

Codon optimized constructs were tested for protein expression using ELISA as previously described by Ghosh et al. ^34^. Briefly, 1 µg of plasmid DNA was used to transfect 1 mL of Gibco^™^Expi293F^™^ cells (Thermo Fisher Scientific) in a 2 mL 96-deepwell plate using FectoPRO^®^ (Polyplus) transfection reagent, per the manufacturer’s guidelines. Twenty-four hours post-transfection, each well was supplemented with 25 µL of 300 mM Valproic Acid (Sigma-Aldrich) and D-(+)-Glucose solution (Sigma-Aldrich) at a 10:9 ratio. Proteins were expressed at 37°C for 4 days post-feeding. On day 5, 150 uL secreted proteins were harvested and biotinylated overnight with 0.25 ug of Biotin Ligase in the presence of 10 mM ATP and 10 mM Mg(OAc)_2_(Sigma-Aldrich) at 4°C.

Expression yields were quantified by ELISA. Biotinylated supernatants (55 µL) were incubated in anti-HIS antibody coated plates. Bound biotinylated proteins were detected using IRDye^®^ 800CW Streptavidin (LI-COR Biosciences). A serial dilution of biotinylated hLOX-1 was used to generate a standard curve. The standard curve was fitted using a four-parameter logistic (4PL) sigmoidal model, plotting concentration (nM) on the x-axis against fluorescence intensity on the y-axis. Biotinylated antigen concentrations were interpolated from the curve, and expression yields of identical proteins were compared across the five codon optimization strategies.

### Large-scale expression and purification of secreted recombinant proteins

Constructs predicted to express at high (>100 nM), medium (50-100 nM) or low (20-10 nM) levels were selected for large scale recombinant protein expression and purification. Gibco^™^ Expi293F^™^ cells (50-200 mL cultures) were transfected with 0.8 µg/mL of plasmid DNA using FectoPRO^®^ (Polyplus) transfection reagent according to the manufacturer’s instructions and incubated at 37°C for 18 – 20 hours. Cultures were then fed with 300 mM Valproic Acid (Sigma-Aldrich) and D-(+)-Glucose solution (Sigma-Aldrich) at a 10:9 ratio, followed by expression at 32°C for 4 days.

Secreted proteins were harvested by centrifugation at 5000 x g and 4°C and purified by gravity-flow nickel affinity chromatography using cOmplete His-Tag purification resin (Roche). A 50% slurry of nickel beads, pre-equilibrated in Binding Buffer (20 mM HEPES pH 8, 250 mM NaCl, 1 mM CaCl_2_, and 1 mM MgCl_2_), was added at the ratio 1 mL beads to 50 mL supernatant and incubated for 1 hour at 4°C. After removal of unbound proteins, the beads were washed twice with 20 mL Binding Buffer containing 20 mM imidazole, and bound proteins were eluted following a 5-minute incubation with 2.6 mL Binding Buffer containing 250–350 mM imidazole.

Eluates with an absorbance greater than 0.3 at 280 nm were further purified by size exclusion chromatography on a HiLoad 16/600 Superdex 200 pg column using an AKTA Pure 25 L (Cytiva). Final protein yields were determined from the absorbance at 280 nm using the calculated extinction coefficients for each protein. Protein identity and molecular weight were confirmed by SDS–PAGE followed by Coomassie staining. Protein monodispersity and post–freeze/thaw stability were assessed by dynamic light scattering (DLS).

## Supplementary Tables

**Table S1.** Epitope tag antibodies production yields using the pTipi2.1 vector. Yields obtained from 100 mL cultures of ExpiHEK cells after cotransfection of the heavy chain (HC) and light chain (LC) plasmids at a 1:2.5 HC/LC ratio. Addgene deposition codes for antibody protein and for the heavy/light chain plasmids.

| Antibodies | Yield (mg/L) | RRIDs | Addgene # for antibody | Addgene # for expression plasmids (HC/LC) |
| --- | --- | --- | --- | --- |
| Anti-Biotin (M33) | 120.0 | AB_3095601 | #218102-rAb | #221349/#221350 |
| Anti-EE (3D5EE_48.K) | 120.0 | AB_3095602 | #218103-rAb | #221351/#221352 |
| anti-GCN4 (C11L34) | 76.9 | AB_3095603 | #218104-rAb | #221353/#221354 |
| anti-Strep (C23.21) | 117.6 | AB_3095604 | #218105-rAb | #221355/#221356 |
| anti-Flag (4E11/M1) | 139.9 | AB_3095605 | #218106-rAb | #221357/#221358 |
| anti-V5 (SV5-Pk1) | 13.8 | AB_3095606 | #218107-rAb | #221359/#221360 |
| anti-ProC (HPC-4) | 99.1 | AB_3095607 | #218108-rAb | #221361/#221362 |
| anti-His (HAP1/5.13.22) | 123.5 | AB_3095608 | #218109-rAb | #221363/#221364 |
| anti-Rho (Rho 1D4) | 159.7 | AB_3095609 | #218110-rAb | #221365/#221366 |

**Table S2.** Codon usage for the human LRP6_26-630 construct for Leucine. The six degenerate Leucine codons are listed together with the usage percentage in the entire human proteome. The native codon scheme is obtained from the translated RNA sequence from transcript 1 from the Refseq database with entry number NM_002336. Codon optimization was performed using proprietary software made available by companies A, B, C & D that provide synthetic genes. The Harmonized scheme was obtained using the Charming tool ^32^, the Skewed only uses the most abundant codon CTG, and the LinearDesign scheme is obtained after optimizing RNA stability ^10^.

| Codon | Human genome usage | Native | Harmonized | CompA | CompB | CompC | CompD | Skewed | LinearDesign |
| --- | --- | --- | --- | --- | --- | --- | --- | --- | --- |
| CTG | 40% | 8 | 6 | 51 | 57 | 58 | 14 | 60 | 33 |
| CTC | 20% | 6 | 16 | 4 | 3 | 2 | 15 | 0 | 22 |
| TTG | 13% | 23 | 7 | 1 | 0 | 0 | 10 | 0 | 2 |
| CTT | 13% | 9 | 13 | 3 | 0 | 0 | 15 | 0 | 3 |
| TTA | 7% | 12 | 10 | 1 | 0 | 0 | 6 | 0 | 0 |
| CTA | 7% | 2 | 8 | 0 | 0 | 0 | 0 | 0 | 0 |

**Table S3.** Sequence sources and identities for the Wnt receptor and carrier glycoprotein constructs considered in this study. Gene names, Refseq and Uniprot identities, construct boundaries and sequence identity for each construct between human and mouse homologue.

| Human gene name | RefSeq ID | UniProt ID | Construct | Murine gene name | Refseq ID | Uniprot ID | Construct | Sequence identity (%) |
| --- | --- | --- | --- | --- | --- | --- | --- | --- |
| LRP4 | NM_002334 | O75096 | 439 - 1351 | Lrp4 | NM_172668 | Q8VI56 | 439 - 1351 | 97 |
| LRP5 | NM_002335 | O75197 | 32 - 643 | Lrp5 | NM_008513 | Q91VN0 | 32 - 642 | 95 |
| LRP5 | NM_002335 | O75197 | 642 - 1254 | Lrp5 | NM_008513 | Q91VN0 | 642 - 1253 | 95 |
| LRP6 | NM_002336 | O75581 | 20 - 630 | Lrp6 | NM_008514 | O88572 | 30 - 630 | 98 |
| LRP6 | NM_002336 | O75581 | 629 - 1244 | Lrp6 | NM_008514 | O88572 | 629 - 1244 | 98 |
| ROR1 | NM_005012 | Q01973 | 30 - 403 | Ror1 | NM_013845 | Q9Z139 | 30 - 403 | 97 |
| ROR2 | NM_004560 | Q01974 | 34 - 402 | Ror2 | NM_013846 | Q9Z138 | 34 - 402 | 94 |
| SFRP1 | NM_003012 | Q8N474 | 32 - 314 | Sfrp1 | NM_013834 | Q8C4U3 | 32 - 314 | 97 |
| SFRP2 | NM_003013 | Q96HF1 | 25 - 295 | Sfrp2 | NM_009144 | P97299 | 25 - 295 | 98 |

**Table S4.**
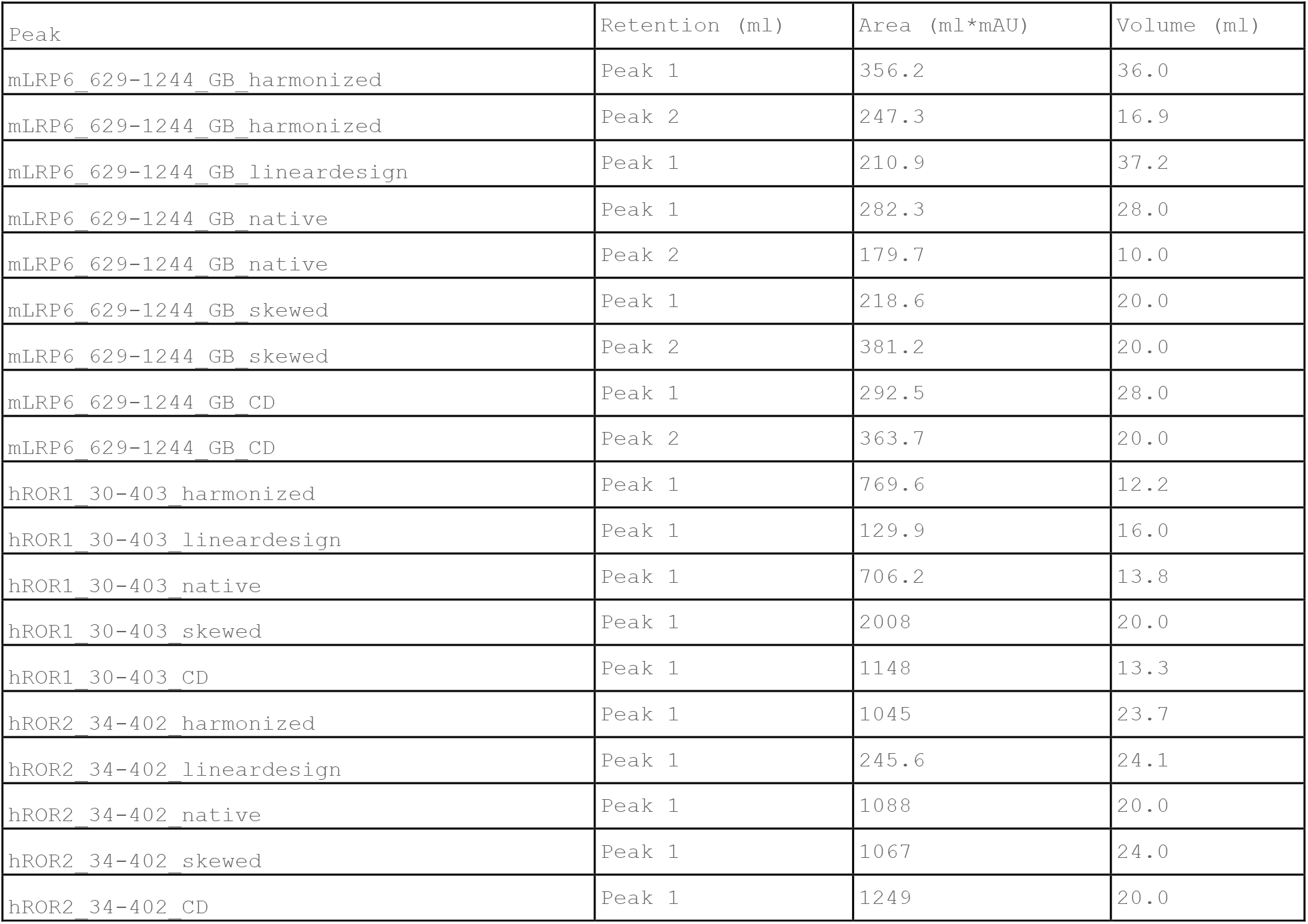
Comparison of protein yield based on the area under the curve in size exclusion chromatography. Protein yields for the human ROR1, ROR2 and murine LRP6 constructs were measured by calculating the area under the curve obtained from the UV detector during size exclusion chromatography.

| Peak | Retention (ml) | Area (ml*mAU) | Volume (ml) |
| --- | --- | --- | --- |
| mLRP6_629-1244_GB_harmonized | Peak 1 | 356.2 | 36.0 |
| mLRP6_629-1244_GB_harmonized | Peak 2 | 247.3 | 16.9 |
| mLRP6_629-1244_GB_lineardesign | Peak 1 | 210.9 | 37.2 |
| mLRP6_629-1244_GB_native | Peak 1 | 282.3 | 28.0 |
| mLRP6_629-1244_GB_native | Peak 2 | 179.7 | 10.0 |
| mLRP6_629-1244_GB_skewed | Peak 1 | 218.6 | 20.0 |
| mLRP6_629-1244_GB_skewed | Peak 2 | 381.2 | 20.0 |
| mLRP6_629-1244_GB_CD | Peak 1 | 292.5 | 28.0 |
| mLRP6_629-1244_GB_CD | Peak 2 | 363.7 | 20.0 |
| hROR1_30-403_harmonized | Peak 1 | 769.6 | 12.2 |
| hROR1_30-403_lineardesign | Peak 1 | 129.9 | 16.0 |
| hROR1_30-403_native | Peak 1 | 706.2 | 13.8 |
| hROR1_30-403_skewed | Peak 1 | 2008 | 20.0 |
| hROR1_30-403_CD | Peak 1 | 1148 | 13.3 |
| hROR2_34-402_harmonized | Peak 1 | 1045 | 23.7 |
| hROR2_34-402_lineardesign | Peak 1 | 245.6 | 24.1 |
| hROR2_34-402_native | Peak 1 | 1088 | 20.0 |
| hROR2_34-402_skewed | Peak 1 | 1067 | 24.0 |
| hROR2_34-402_CD | Peak 1 | 1249 | 20.0 |

**Table S5.** Golden Gate Part Types.

| Part Number | Part Type | Upstream Overhang | Downstream Overhang |
| --- | --- | --- | --- |
| 0 | Part Entry Vector | AACG | GCTG |
| 1 | Backbone | AGTG | CTGA |
| 2 | N-Term | AGTG | CTCC |
| 3 | CDS | CTCC | AGGA |
| 4 | C-Term | AGGA | CTGA |

## Supplementary Figures

**Figure S1.**
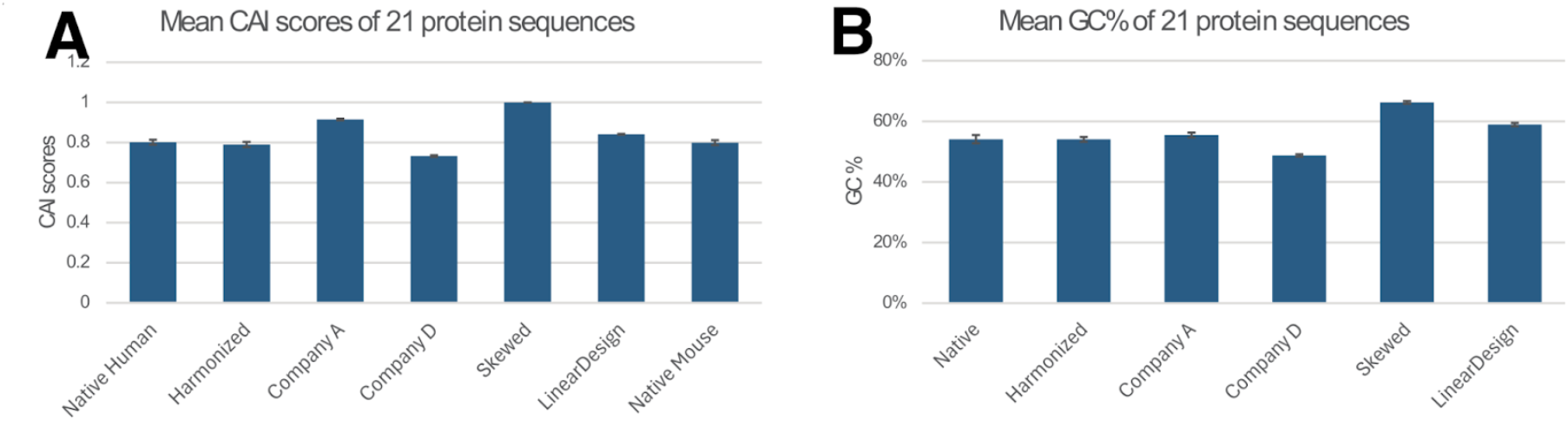
Codon adaptation index (CAI) and GC content for each codon scheme for the 18 constructs.

**Figure S2.**
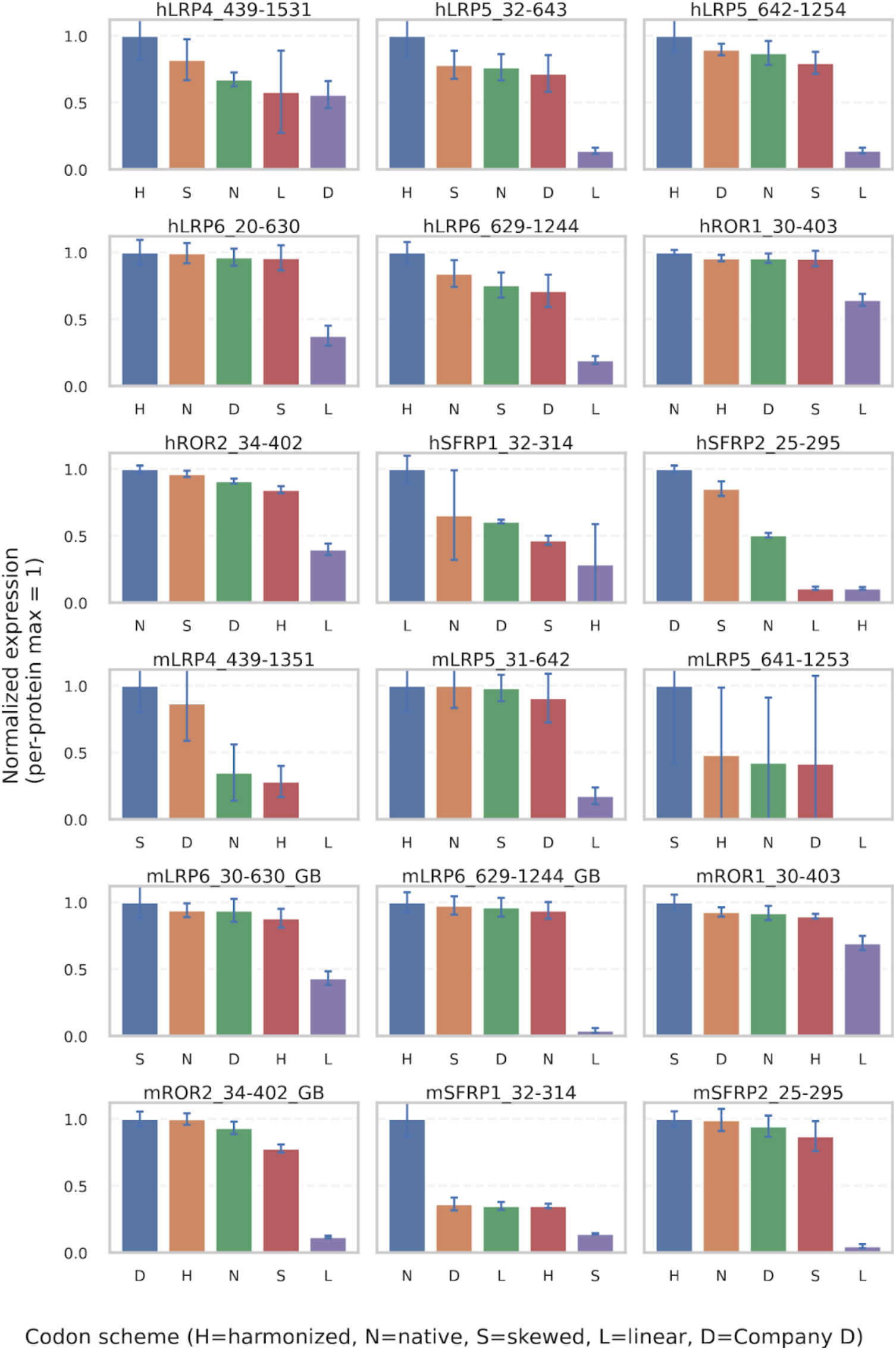
Normalized expression levels are shown for 18 individual proteins expressed using five different codon schemes (harmonized, linear design, native, skewed, and Company D). For each protein, fluorescence values were first averaged across triplicate measurements and then normalized to the highest-expressing codon scheme for that protein (set to 1.0), allowing direct comparison of relative performance within each protein. Bars represent normalized expression, and error bars indicate the normalized standard error derived from the triplicate measurements. Within each panel, codon schemes are ordered from highest to lowest normalized expression.

